# A cardiotonic steroid multiplex method using ultra-high-performance liquid chromatography-tandem mass spectrometry

**DOI:** 10.1101/2023.10.07.561354

**Authors:** Lorna C Gilligan, Daniyah Alamrani, Dannie Fobian, Dipak Kotecha, Paulus Kirchhof, Wiebke Arlt, Angela E Taylor, Davor Pavlovic

## Abstract

**Background and Aims:** Digoxin, a cardiotonic steroid (CTS), is commonly prescribed for patients with atrial fibrillation and heart failure. Endogenous CTS have been implicated in cardiovascular disease pathogenesis and can interact with digoxin. We developed an ultra-high-performance liquid chromatography-tandem mass spectrometry (UHPLC-MS/MS) assay for the quantification of eleven CTS.

**Materials and Methods:** Isotopically labelled internal standards were added to samples, followed by protein precipitation and solid-phase extraction. Steroids were separated using an Acquity uPLC chromatography system with a Waters CORTECS T3 column (1.6 μm 2.1x50 mm) and quantification performed on a Waters TQ-XS mass spectrometer using electrospray ionisation in positive ion mode. Separation used a methanol/ water elution system containing 0.1% formic acid and post-column infusion of lithium chloride.

**Results:** Run time was 13.5 minutes. Lower limits of quantification ranged from 0.025 to 0.1 ng/ml. Serum recovery ranged from 40.0-98.6% with matrix effects from -22.9% to 7.6%. Plasma recovery ranged from 27.4-83.9% and matrix effects were -27.6-39.1%. Accuracy and precision at three concentrations were within ideal range (<15%) for seven CTS and <20% for the others.

**Conclusion:** This validated UHPLC-MS/MS method provides a comprehensive assessment profiling 11 cardiotonic steroids, offering a sensitive and specific tool for clinical and pre-clinical investigations.

**Highlights:**

- Mass spectrometry method simultaneously quantifies multiple cardiotonic steroids
- Accurate and specific measurement of clinically relevant digoxin concentration
- Method validated for measurement of cardiotonic steroids in serum and plasma
- Post-column infusion of lithium chloride substantially improves sensitivity

**Research funding:** This work was funded by the British Heart Foundation (PG/17/55/33087, FS/PhD/22/29309, FS/19/12/34204, RG/17/15/33106 to DP, Accelerator Award AA/18/2/34218 to Institute of Cardiovascular Sciences), Wellcome Trust (Seed Award Grant 109604/Z/15/Z to DP) and Department of Clinical Laboratory Sciences, Faculty of Applied medical Sciences, University of Hail. PK was partially supported by European Union AFFECT-AF (grant agreement 847770), and MAESTRIA (grant agreement 965286), British Heart Foundation (PG/17/30/32961; PG/20/22/35093; AA/18/2/34218), German Centre for Cardiovascular Research supported by the German Ministry of Education and Research (DZHK), Deutsche Forschungsgemeinschaft (Ki 509167694), and Leducq Foundation. The funding organization(s) played no role in the study design; in the collection, analysis, and interpretation of data; in the writing of the report; or in the decision to submit the report for publication.

**Financial disclosures:** Prof. Kotecha reports grants from the National Institute for Health Research (NIHR CDF-2015-08-074 RATE-AF; NIHR130280 DaRe2THINK; NIHR132974 D2T-NeuroVascular; NIHR203326 Biomedical Research Centre), the British Heart Foundation (PG/17/55/33087, AA/18/2/34218 and FS/CDRF/21/21032), the EU/EFPIA Innovative Medicines Initiative (BigData@Heart 116074), EU Horizon (HYPERMARKER 101095480), UK National Health Service -Data for R&D-Subnational Secure Data Environment programme, UK Dept. for Business, Energy & Industrial Strategy Regulators Pioneer Fund, the Cook & Wolstenholme Charitable Trust, and the European Society of Cardiology supported by educational grants from Boehringer Ingelheim/BMS-Pfizer Alliance/Bayer/Daiichi Sankyo/Boston Scientific, the NIHR/University of Oxford Biomedical Research Centre and British Heart Foundation/University of Birmingham Accelerator Award (STEEER-AF). In addition, he has received research grants and advisory board fees from Bayer, Amomed and Protherics Medicines Development; all outside the submitted work.

PK received research support for basic, translational, and clinical research projects from European Union, British Heart Foundation, Leducq Foundation, Medical Research Council (UK), the Deutsche Forschungsgemeinschaft (DFG) and German Centre for Cardiovascular Research, from several drug and device companies active in atrial fibrillation, and has received honoraria from several such companies in the past, but not in the last three years. PK is listed as inventor on two issued patents held by University of Hamburg (Atrial Fibrillation Therapy WO 2015140571, Markers for Atrial Fibrillation WO 2016012783).

**Graphical Abstract:** 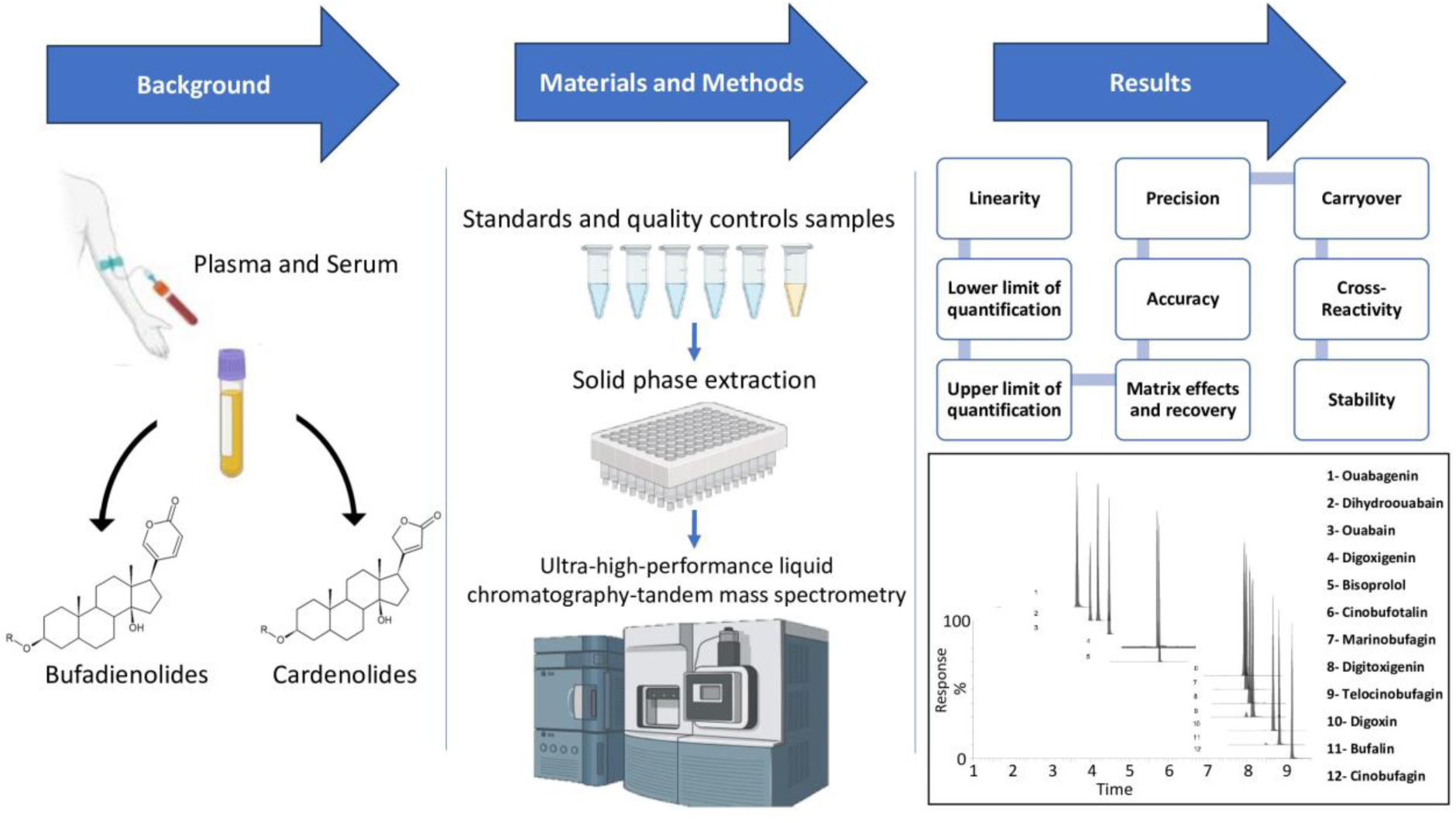

## 1.0 Introduction

Cardiotonic steroids (CTS) are a class of biologically active small molecules. The best-known CTS, digoxin, is used as a rate-controlling and positive inotropic agent in the treatment of patients with heart failure and atrial fibrillation.

CTS structurally consist of a steroid backbone with either a 5-membered unsaturated (cardenolides) or a 6-membered doubly unsaturated (bufadienolides) lactone ring at position 17. Cardenolides have a hydroxyl at position 14 and some bufadienolides have an epoxide at position 14-15. The steroid core can also be glycosylated at position 3 [1] (**Figure 1**). CTS can bind and inhibit the ubiquitous NA+/K+-ATPase, raising intracellular sodium, which leads to reduced activity of the sodium-calcium exchanger and increases intracellular calcium. This can have a positive inotropic effect leading to increased cardiac contractility. CTS also increase vagal tone and thereby reduces heart rate.

**Figure 1.**
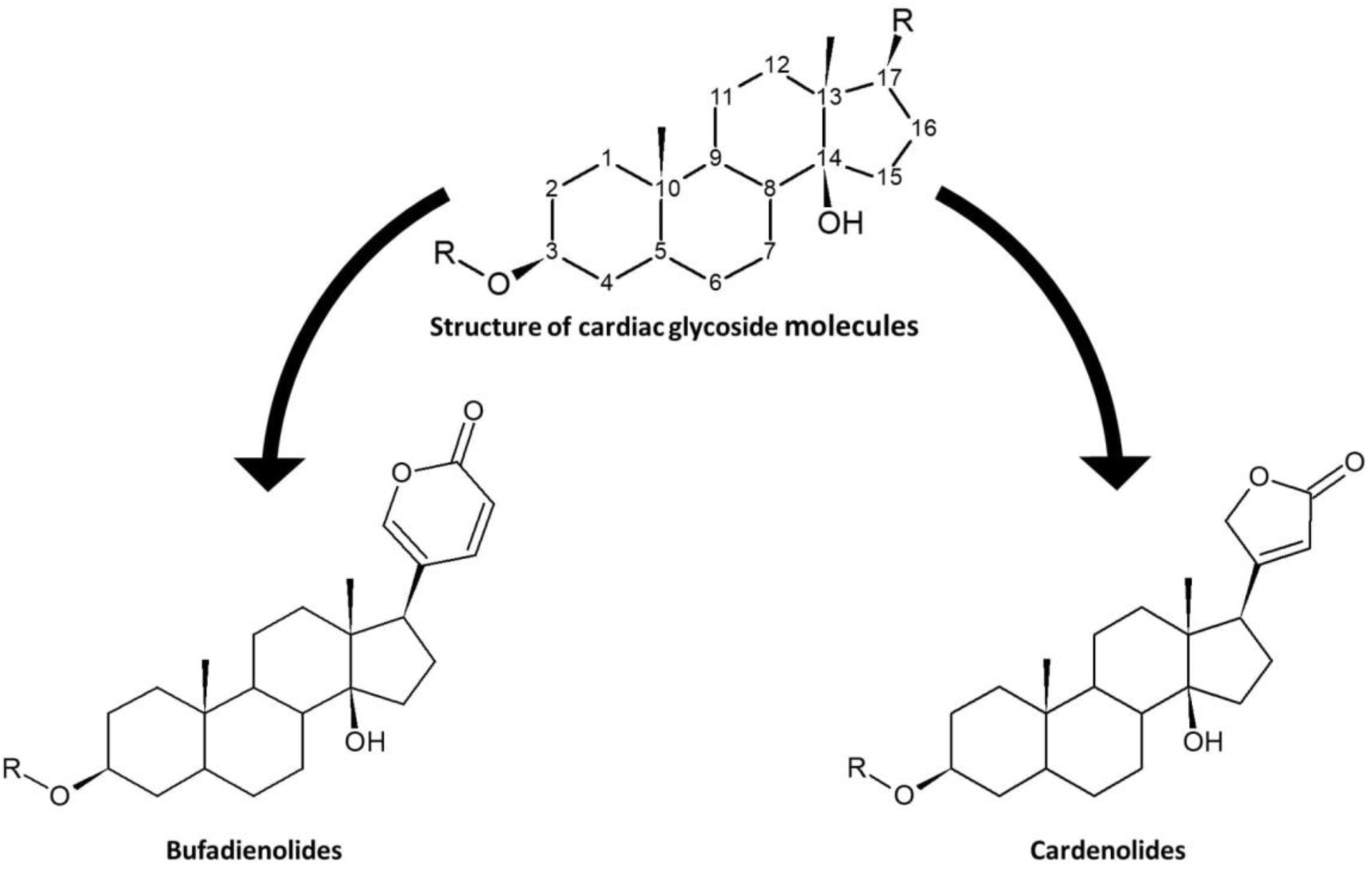
Structure of cardenolides and bufadienolides. Cardiotonic steroids have the four-ring steroid backbone, at position 17 (R) cardenolides have a five-membered lactone ring, and bufadienolides a six-membered lactone ring. RO is a sugar (glycoside) which is different for each CTS.

Endogenous cardiotonic steroids were first detected in human circulation after a series of studies conducted by Hamlyn and colleagues [2,3]. Since then, different immunoassay techniques have been developed to measure CTS in humans and animals [4–7]. Cardenolides, such as ouabain, have been detected and measured in the blood of healthy people and those with congestive heart failure [4], hypertension [5], acute myocardial infarction [6], and renal disease [7]. Bufadienolides have been detected in blood and urine from patients with sodium loading and plasma volume expansion conditions such as chronic renal failure [8], congestive heart failure [9], and pre-eclampsia [10]. CTS furthermore have been isolated from mammalian adrenal tissue, which is thought to be the site of their synthesis [11,12]. CTS may also be produced in the heart [13], placenta [14], and hypothalamus [15]. Their synthetic pathway from cholesterol has yet to be elucidated, though many factors have been identified that influence their metabolism such as adrenocorticotropic hormone [11,16], angiotensin II [11], volume expansion [17], increase in physical activity [18], and hypoxia [19].

Digoxin, derived from the purple foxglove plant, Digitalis purpurea, has been prescribed to treat heart failure for over 200 years [1,20,21]. Digoxin is routinely measured clinically by immunoassay. Low doses of digoxin are often well-tolerated [22,23] while higher concentrations lead to toxicity. Toxicity events can occur in some patients even within the narrow therapeutic range. It has been suggested that CTS may interfere with digoxin and thus contribute to this toxicity, however, the presence of endogenous CTS and their physiological and pathological roles remain controversial. Using enzyme-linked immunoassay (ELISA) based methods, nanomolar concentrations of the CTS, ouabain, [24,25], digoxin [26], marinobufagenin (MBG) [10], telocinobufagin (TBG) [27], and bufalin [28] have been reported in human serum.

Several methods utilising liquid chromatography coupled with mass spectrometry (LC-MS) have identified CTS such as ouabain [25], digoxin [25], MBG [29], TBG [27], and bufalin [30]. However, a study by Baecher et al. was unable to detect ouabain in biological samples even with a sensitive, well-validated mass spectrometry-based assay [31]. Similarly, Lewis et al. [32] were able to detect ouabain with ELISA alone but failed to identify ouabain using ELISA combined with high-pressure liquid chromatography (HPLC) fractionation. The variation between laboratories is in part due to the variety of extraction procedures and analytical technologies used for quantification, as well as significant differences in the patient populations investigated. These differing results have led to heated debate within the field [33–37]. A reliable assay quantifying several cardiotonic steroids at physiological concentrations would be of benefit to this field.

We aimed to develop an assay capable of accurately identifying and quantifying several CTS, enabling further understanding of the roles of each CTS and the metabolic pathways involved. We aimed to develop, optimise, and validate a method that combined a single extraction method with liquid chromatography-tandem mass spectrometry (LC-MS/MS). LC-MS/MS has increased specificity compared to immunoassay as cross-reactivity from other CTS is no longer problematic, permitting its use as a profiling tool for CTS in biological samples.

The steroids measured in this assay should cover a range of cardenolides and bufadienolides. We optimised a method for the detection and quantification of cardenolides ouabain, dihydroouabain (DHO), ouabagenin, digoxigenin, digitoxigenin, and digoxin, and bufadienolides bufalin, cinobufotalin, cinobufagin, marinobufagin (MBG), and telocinobufagin (TBG). A final requirement of the method was sufficient sensitivity to cover the expected concentration ranges in human serum and plasma. There is limited information on the expected concentration ranges of all CTS in humans, therefore we investigated a similar concentration range based on the widely studied CTS, ouabain of <0.4 to more than 8.7 nM (∼0.23 to 5 ng/mL) [4,38].

## 2.0 Materials and Methods

### 2.1 Preparation of standards and quality controls

Standards of all analytes were purchased as powders (**Table 1**). Individual stock solutions at 1 mg/mL were prepared in UHPLC grade methanol (Biosolve), or for isotopically labelled internal standards, to avoid hydrogen-deuterium exchange, in deuterated methanol (Sigma-Aldrich), all were stored at -80 °C. Each steroid was infused into the LC-MS/MS in combination with LiCl and cone voltages, collision energies and mass transitions optimised. The CTS 1 mg/ml solutions were combined, and a calibration series was prepared by spiking into a surrogate sample matrix (phosphate buffered saline pH 7.4 supplemented with 0.1% (w/v) bovine serum albumin (Sigma-Aldrich)) at the following concentrations, 0, 0.025, 0.05, 0.1, 0.5, 1, 2.5, 5, 10, 12.5, and 25 ng/mL. Where an isotope labelled internal standard was available this was used for quantification, where this was not available a surrogate internal standard was used. Ouabagenin and DHO used ouabain-d3, cinobufotalin, marinobufagenin, and telocinobufagin used cinobufagine-d3, bufalin, and bisoprolol used digoxigenin-d3. PBS spiked with BSA was used as a surrogate matrix for serum and plasma, as it is routinely used in clinical biochemistry and research laboratories. An internal standard stock solution containing 50 ng/mL of each internal standard was also prepared in UHPLC grade methanol (Biosolve) and stored at -20 °C.

**Table 1.** Cardiotonic Steroid Mass Spectrometry Parameters. Cardiotonic steroid names, molecular formulas, molecular weights (atomic mass units), mass transitions (multiple reaction monitoring mode MRM), retention times (Rt), cone voltages, collision energies, and suppliers. Surrogate internal standards were used for ouabagenin and DHO (ouabain-d3), cinobufotalin marinobufagenin and telocinobufagin (cinobufagine-d3), bufalin and bisoprolol (digoxigenin- d3).

| Analyte<br>(Molecular<br>formula) | Mol<br>weight<br>(amu) | Mass<br>Transitions<br>Quantifier<br>Qualifier | Rt<br>(min) | Cone<br>Voltage<br>(V) | Collison<br>Energy<br>(eV) | Supplier |
| --- | --- | --- | --- | --- | --- | --- |
| Ouabagenin<br>C<sub>23</sub>H<sub>34</sub>O<sub>8</sub> | 438 | 445.2 > 379.1<br>445.2 > 361.1 | 3.7 | 58<br>58 | 38<br>42 | Sigma Aldrich, UK |
| Dihydroouabain<br>C<sub>29</sub>H<sub>46</sub>O<sub>12</sub> | 586 | 593.2 > 447.2<br>593.2 > 381.1 | 4.2 | 100<br>100 | 48<br>54 | Santa Cruz<br>Biotech, DE |
| Ouabain<br>C<sub>29</sub>H<sub>44</sub>O<sub>12</sub> | 584 | 591.2 > 445.1<br>591.2 > 379.1 | 4.5 | 100<br>100 | 48<br>54 | Sigma Aldrich, UK |
| Ouabain-d3 | 588 | 594.2 > 448.1<br>594.2 > 382.1 | 4.5 | 86<br>86 | 48<br>56 | Toronto Research<br>Chemicals, CA |
| Digoxigenin<br>C<sub>23</sub>H<sub>34</sub>O<sub>5</sub> | 391 | 429.1 > 397.1<br>429.1 > 379.1 | 5.7 | 4<br>4 | 12<br>28 | Sigma Aldrich, UK |
| Digoxigenin-d3 | 394 | 432.1 > 400.2<br>432.1 > 382.1 | 5.7 | 28<br>28 | 20<br>32 | Toronto Research<br>Chemicals, CA |
| *Bisoprolol<br>C<sub>18</sub>H<sub>31</sub>NO<sub>4</sub> | 325 | 326.1 > 115.93<br>326.1 > 73.86 | 5.8 | 80<br>80 | 22<br>18 | Sigma Aldrich, UK |
| Marinobufagenin<br>C<sub>24</sub>H<sub>32</sub>O<sub>5</sub> | 400 | 439.2 > 407.2<br>439.2 > 371.2 | 7.9 | 42<br>42 | 14<br>34 | BOC Sciences,<br>USA |
| Cinobufotalin<br>C<sub>26</sub>H<sub>34</sub>O<sub>7</sub> | 458 | 497.2 > 465.2<br>497.2 > 447.2 | 7.9 | 44<br>44 | 8<br>30 | Sigma Aldrich, UK |
| Digitoxigenin<br>C<sub>23</sub>H<sub>34</sub>O<sub>4</sub> | 374 | 413.2 > 381.1<br>413.2 > 363.1 | 8.0 | 6<br>6 | 12<br>30 | Sigma Aldrich, UK |
| Digitoxigenin-d3 | 377 | 416.2 > 384.1<br>416.2 > 366.1 | 8.0 | 28<br>28 | 18<br>34 | Toronto Research<br>Chemicals, CA |
| Telocinobufagin<br>C<sub>24</sub>H<sub>34</sub>O<sub>5</sub> | 402 | 441.2 > 409.2<br>441.2 > 355.2 | 8.1 | 34<br>34 | 10<br>40 | BOC Sciences,<br>USA |
| Digoxin<br>C<sub>41</sub>H<sub>64</sub>O<sub>14</sub> | 780 | 787.4 > 267.1<br>787.4 > 137.0 | 8.7 | 100<br>100 | 48<br>58 | Sigma Aldrich, UK |
| Digoxin-d3 | 784 | 790.4 > 267.1<br>790.4 > 137.0 | 8.7 | 94<br>94 | 48<br>60 | Toronto Research<br>Chemicals, CA |
| Bufalin<br>C<sub>24</sub>H<sub>34</sub>O<sub>4</sub> | 386 | 425.2 > 393.2<br>425.2 > 375.2 | 8.8 | 46<br>46 | 16<br>32 | Sigma Aldrich, UK |
| Cinobufagin<br>C<sub>26</sub>H<sub>34</sub>O<sub>6</sub> | 442 | 481.2 > 449.2<br>481.2 > 389.2 | 9.1 | 40<br>40 | 10<br>32 | Sigma Aldrich, UK |
| Cinobufagine-d3 | 445 | 484.2 > 452.2<br>484.2 > 389.1 | 9.1 | 26<br>26 | 22<br>34 | Toronto Research<br>Chemicals, CA |

\* Not a cardiotonic steroid, included in assay as often prescribed for cardiac related conditions such as AF.

To continually monitor batch accuracy and precision of the method two quality control sample types were prepared. Quality control (QC) and biological QC were prepared independently to the calibrators. First to measure accuracy QCs were prepared by spiking the surrogate serum matrix at three concentrations, representing the approximate literature quoted concentration ranges for ouabain in human serum (0.3, 1.5, and 15 ng/mL). Each concentration was extracted with each batch, with a batch acceptability criteria of a bias of +/- 15% from the expected nominal concentrations. Secondly to monitor batch precision, pooled adult male human serum (Sigma-Aldrich) or pooled human plasma (six adults, three males, and three females) were spiked with a mix of all CTS at 2 ng/ml and stored at -20 °C. These samples were extracted in triplicate with each batch, with a batch acceptability criteria of precision (% variation) or <20%. These quality control samples were also used during the validation procedure.

### 2.2 Sample preparation

Calibrators, QCs, and biological QCs were extracted with each batch. 500 µL of sample, calibrator, QC, or biological QC were transferred into a 2mL 96-well plate (Porvair Sciences) and 10 µL of the internal standards mixture (final concentration of 1 ng/ml) containing all stable isotope labelled internal standards listed in **Table 1** were added. 100 μL of ZnSO_4_ (Sigma Aldrich, UK) was added to precipitate the proteins and the samples were vortexed for 5 minutes on a Fisherbrand multi-tube vortex (Fisher, UK). 400 μL of water (Fisher) was added before centrifugation at g x 976 (3000 rpm) for 10 minutes on an Eppendorf 5810 centrifuge. Using a positive pressure manifold (Biotage), the SPE plate (Thermo SOLA SPE 10 mg/ 2ml HRP) was activated with 500 μL of UHPLC grade methanol (Biosolve) and equilibrated with 500 μL of UHPLC grade water (Fisher). The top 800 μL of the sample was passed through the SPE avoiding transfer of the protein precipitate. Subsequently, the SPE plate was washed with UHPLC grade water. Finally, 250 μL x 2 of UHPLC grade methanol was passed through the cartridge and collected in a 96-well plate containing 700 µL glass inserts (Randox). The eluate was dried under a stream of nitrogen. The dried extract was reconstituted in 200 µL of 10% (v/v) UHPLC grade methanol (Biosolve) in UHPLC grade water.

### 2.3 Ultra-high performance liquid chromatography

The liquid chromatography separation was optimised using a combination of automated MUSCLE (Multi-objective Unbiased optimisation of Spectrometry via Closed Loop Experimentation) software and traditional method development techniques [39]. Chromatography was performed on an ACQUITY ultra-performance liquid chromatography system (UPLC; Waters) using a CORTECS T3 1.6 μm 2.1x50 mm column at 35 °C. The injection volume was 20 µL. The CTS were chromatographically resolved using a novel convex gradient profile of A water (0.1% formic acid) (Fisher chemical) and B methanol (0.1% formic acid) (Biosolve) with post-column infusion of lithium chloride (LiCl). To separate the analytes, a flow rate of 0.5 mL/min was applied with a linear gradient from 10% to 30% B over five minutes, followed by a convexly curved gradient from 45% to 95% B over 6.5 minutes. The column was then equilibrated for 1.5 minutes at starting conditions prior to the injection of the next sample (**Figure 2**). The autosampler was maintained at 10 °C.

**Figure 2.**
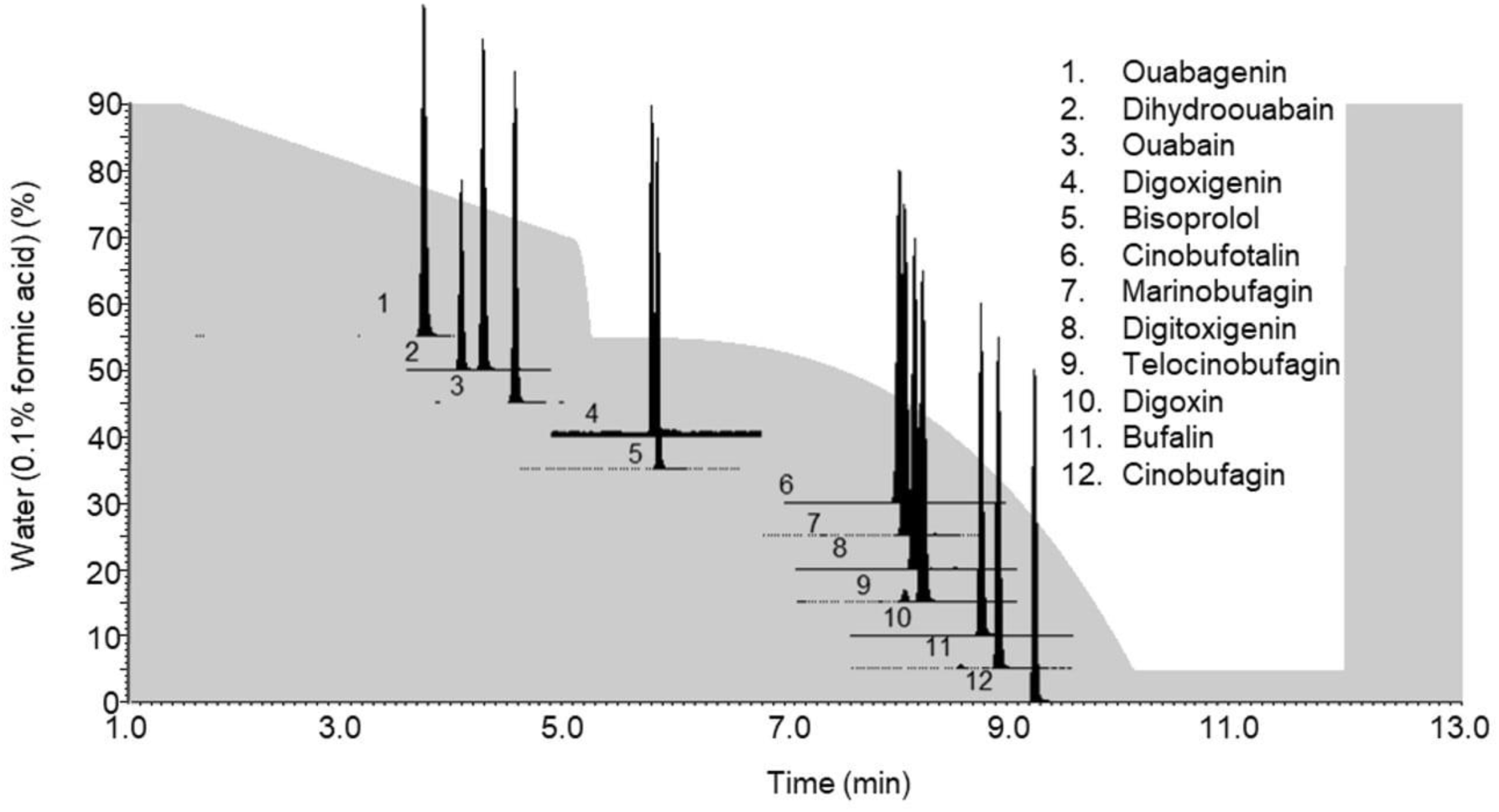
Chromatographic separation of cardiotonic steroids. Gradient profile demonstrating the change in percentage water (0.1% formic acid) over time (grey shaded area), overlaid with the chromatographic peaks of the 11 CTS and bisop

### 2.4 Tandem mass spectrometry

The UHPLC eluate was combined post column with 1 mM LiCl (in 95% UHPLC grade methanol and 5% UHPLC grade water at a constant flow rate of 5 μL/min) and directed into a XEVO^TM^ TQ-XS mass spectrometer (Waters, UK) using electrospray ionisation in positive ion mode. The capillary voltage was maintained at 0.6 kV, the source temperature was 150 °C, the desolvation temperature was 500 °C, the gas flow was 1000 L/h, and the cone gas was 150 L/h. MassLynx© 4.2 software (Waters, UK) was used for system control. CTS were quantified based on their lithium adducts using two mass transitions, (a quantifier and a qualifier). The mass transitions and respective cone voltages and collision energies are summarised in **Table 1**. Dwell time was automatically calculated by the MassLynx© software for each CTS with a minimum of 12 analytical points across each peak for quantitative analysis. TargetLynx software was used for data processing and quantification. For quantitation, peak area ratios of analyte to internal standard were plotted against the nominal concentrations of the calibrators, and 1/x weighting was applied to produce the calibration curves.

### 2.5 Method Validation

Validation was performed following protocols developed in our lab and from published guidelines [40–43].

#### 2.5.1 Linearity

Three calibration series were prepared independently and processed on different days. For each analyte, the ratio of the analyte peak area to the internal standard peak area was plotted against the nominal concentrations of the standard. Calibration curves were accepted as linear if the correlation coefficient of the linear regression (R^2^) was ≥0.99 and each data point above the lower limit of quantitation (LLOQ) and below the upper limit of quantitation (ULOQ) had an accuracy (bias) of <±15% compared to the theoretical concentration.

#### 2.5.2 Lower limit of quantification

The lower limit of quantification was defined as the lowest concentration for which 10 replicate samples of CTS spiked surrogate matrix (PBS with 0.1%BSA) could be quantified with an intra-assay precision measured as the percentage relative standard deviation (RSD %) of <20% and an accuracy (bias) of not greater than ±20%.

#### 2.5.3 Upper limit of quantification

The upper limit of quantification was defined as the highest concentration for which 10 replicate samples of CTS spiked surrogate matrix could be quantified with an intra-assay precision RSD% <15% and with an accuracy (bias) not greater than ±15%.

#### 2.5.4 Precision

The surrogate matrix was spiked with all CTS analytes at three different concentrations (0.3, 1.5, and 15 ng/mL). These samples were extracted and quantified 10 times in the same batch to evaluate intra-assay precision. A relative standard deviation (RSD %) ≤15% was considered acceptable.

Pooled serum and pooled plasma samples were spiked with 2 ng/mL of all analytes and extracted and quantified 10 times on different days to assess inter-assay (between-run) and intra-assay (within-run) precision. A relative standard deviation (RSD %) ≤15% was considered acceptable.

#### 2.5.5 Accuracy

Ten samples of the surrogate matrix were spiked with all CTS at three different concentrations (0.3, 1.5, and 15 ng/mL). Once quantified relative to the calibration series, the percentage bias of the observed concentration to the nominal concentration between ±15% was deemed acceptable.

#### 2.5.6 Matrix effects and recovery

Matrix effects can either lead to a decrease in response (ion suppression) or an increase in response (ion enhancement) of the analytes of interest. These effects can be due to contaminants in the matrix altering the ionisation of the analyte, or due to competition for ionisation from co-eluting compounds.

Matrix effects and extraction efficiency were determined as previously described [44]. Six pooled serum (Sigma-Aldrich) and six plasma samples from healthy volunteers were spiked with a mix of all CTS at 2 ng/mL before extraction (pre-extraction) and after extraction (post-extraction). Additionally, the reconstitution solvent (10% (v/v) UHPLC grade methanol (Biosolve) in UHPLC grade water) was spiked (no extraction). Internal standards were added to all samples. Matrix effects (%) were calculated using the equation *((post-extraction - no extraction)/no extraction) x 100*. Recovery (%) (extraction efficiency) was calculated by comparing pre- to post-extraction samples and converted to a percentage using the equation *(pre-extraction/post-extraction) x 100*. Matrix effects between -20% and 20% were considered acceptable, with those between -15% to 15% ideal. Recovery between 80% and 120% was desirable, although lower recoveries are acceptable for a profiling method if the extraction is reproducible.

#### 2.5.7 Carryover

Carryover was assessed after injection of the top calibrant and a sample ten times the concentration of the top calibrant (25 and 250 ng/mL). Percentage carryover from the samples was calculated by direct comparison of the peak area in the concentrated sample to the subsequently injected blank sample. A carryover of <2% was acceptable.

#### 2.5.8 Cross-Reactivity

The beta-blocker bisoprolol was added to the method to rule out cross-reactivity with CTS as it has a comparable molecular weight and is commonly prescribed in patients with cardiovascular disease [22].

#### 2.5.9 Stability

Stability was assessed at three concentrations (0.3, 1.5, and 15 ng/ml) prepared in surrogate matrix, which were aliquoted and stored at -20°C. Different aliquots were extracted after addition of internal standards eight times over a three-month period. Accuracy expressed as percentage bias from theoretical concentrations and precision expressed as percentage variance was determined in all experiments. CTS could be described as stable if accuracy and precision demonstrated reproducible data over this time.

## 3.0 Results

The CTS included in this method have a range of polarities. This broad polarity range can be challenging for a single extraction and chromatographic method. Several solid-phase extraction (SPE) and column chemistries were tested. Thermo Scientific SOLA SPE plates HRP provided maximum recovery across all analytes, and a CORTECS T3 1.6 μm 2.1x50 mm column produced optimal chromatographic resolution of the CTS.

### 3.1 Chromatography and Mass Spectrometry Optimisation

Two mass transitions were optimised for each CTS (**Table 1**). Analytes were separated using two sequential gradients; one linear, followed by one convex. The following the wash and re- conditioning steps resulted in a total run time injection-to-injection of 13.5 minutes. All CTS and bisoprolol were baseline resolved or mass separated using this chromatographic method, with no cross-reactivity (**Figure 2**).

#### 3.1.1 Optimisation of mobile phase additives

Lithium adducts have been shown to improve the signal intensity of ouabain [34] but to our knowledge have not been used in a multi-analyte CTS profiling method. Analytes were directly infused with LiCl to form lithium adducts. Stable lithiated parent and daughter ions were obtained, with quantitative mass transitions from the largest peak areas selected for the method. LiCl was originally added to the aqueous mobile phase and concentration was titrated to achieve the highest peak area for each of the CTS without any ion suppression. However, the addition of LiCl to the water phase pre-column caused an unacceptable increase in system backpressure, affecting chromatography and blocking the column after a few hundred injections. Therefore, post-column infusion was used as described by Baecher et al. [31]. For the XEVO^TM^ TQ-XS mass spectrometer (Waters, UK), a minimum post-column infusion flow rate of 5 µL/min was required to maintain a stable flow of LiCl into the source. 1 mM LiCl infused post-column achieved stable lithium adducts. The stability of the adduct formation was determined following 20 injections of a 1 ng mixture of all CTS, demonstrating peak area variation for each of <10%. The formation of Li adducts improved the signal for all CTS compared to formic acid alone. Absolute peak areas increased from 235 to 5799% after the formation of Li-adducts (**Supplementary Figure 1**). All analytes demonstrated unique molecular ions and mass transitions (**Table 1**). Bisoprolol did not form a Li adduct and the signal was reduced by half with the inclusion of LiCl. However, due to the high ionisation efficiency of bisoprolol quantification was still possible. 0.1% formic acid was added to the aqueous mobile phase to reduce the risk of bacterial contamination in the water.

### 3.2 Validation of the analytical method

The analytical performance of the optimised assay was evaluated.

#### 3.2.1 Method Linearity

Calibration curves for all analytes were linear between the LLOQ and ULOQ, with R^2^≥0.99 and with acceptable deviations from the nominal concentrations. Lower limits of quantification ranged between 0.025 ng/mL and 0.1 ng/mL for all analytes. The chromatograms shown in **Supplementary Figure 2** demonstrate the peak of each CTS (quantifier mass transition MRM) at the calculated LLOQ for each CTS. Also shown is the corresponding MRM in a blank sample (C0, calibration matrix with internal standards, but no CTS added). 25 ng/mL was the upper limit of quantitation for all CTS, demonstrating an accuracy ranging from -14.8% to - 8.6% and a precision of <10.1% (**Table 2**).

**Table 2:** Accuracy and precision across the calibration range for 11 CTS and bisoprolol. Accuracy (% bias) and precision (% RSD) at the lower (LLOQ) and upper (ULOQ) limits of quantification and at low, medium, and high spiked surrogate matrix. Accuracy (% bias) was determined from ten independently prepared and extracted samples and precision (RSD%) from ten extractions from a single sample.

| Steroid | LLOQ |  |  | ULOQ |  |  | Accuracy (Bias %) |  |  | Precision (RSD %) |  |  |
| --- | --- | --- | --- | --- | --- | --- | --- | --- | --- | --- | --- | --- |
|  | Conc<br>ng/mL | Acc.<br>(Bias %) | Pre.<br>(RSD %) | Conc<br>ng/mL | Acc.<br>(Bias %) | Pre.<br>(RSD %) | Low<br>0.3<br>ng/mL | Med<br>1.5<br>ng/mL | High<br>15<br>ng/mL | Low<br>0.3<br>ng/mL | Med<br>1.5<br>ng/mL | High<br>15<br>ng/mL |
| Ouabagenin | 0.1 | 9.5 | 4.9 | 25 | -9.8 | 8.0 | 3.9 | 10.7 | -1.5 | 7.9 | 5.9 | 7.3 |
| Dihydroouabain | 0.05 | 14.0 | 11.5 | 25 | -12.2 | 10.0 | 4.3 | 9.7 | 1.5 | 6.4 | 5.2 | 5.8 |
| Ouabain | 0.025 | 2.4 | 3.8 | 25 | -13.7 | 8.8 | 5.5 | 6.8 | -1.8 | 7.5 | 4.7 | 3.3 |
| Digoxigenin | 0.05 | 10.4 | 19 | 25 | -11.2 | 9.5 | 16.3 | 4.1 | -5.7 | 8.3 | 5.9 | 3.4 |
| *Bisoprolol | 0.025 | 5.6 | 10.7 | 25 | -6.4 | 10.1 | 8.9 | 2.4 | -1.7 | 15.5 | 9.7 | 6.0 |
| Marinobufagenin | 0.025 | 13.8 | 12.5 | 25 | -13.5 | 8.9 | 11.7 | 11.0 | -14.0 | 9.5 | 15.5 | 6.6 |
| Cinobufotalin | 0.025 | 13.8 | 12.8 | 25 | -14.0 | 9.8 | 13.5 | 12.7 | -10.9 | 8.2 | 15.5 | 6.4 |
| Digitoxigenin | 0.025 | 13.2 | 2.9 | 25 | -8.6 | 9.8 | 6.2 | 4.7 | -1.0 | 13.2 | 6.0 | 4.6 |
| Telocinobufagin | 0.05 | 15.8 | 12.2 | 25 | -14.8 | 8.9 | 13.4 | 13.7 | -13.5 | 11.9 | 20.0 | 5.7 |
| Digoxin | 0.025 | 12.3 | 4.9 | 25 | -12.0 | 9.2 | -0.5 | -3.7 | -10.9 | 6.4 | 6.6 | 4.2 |
| Bufalin | 0.025 | 12.0 | 8.1 | 25 | -11.8 | 9.4 | 12.5 | 11.6 | -3.7 | 6.1 | 12.1 | 5.6 |
| Cinobufagin | 0.025 | 4.4 | 16.3 | 25 | -10.9 | 9.0 | 10.7 | 3.7 | -9.3 | 8.5 | 7.4 | 3.6 |

\* Not a cardiotonic steroid, included in assay as often prescribed for cardiac related conditions such as AF.

#### 3.2.2 Accuracy and Intra and Inter-assay Precision

Accuracy assessed at the three concentrations (0.3, 1.5, and 15 ng/mL) calculated as % bias from the nominal concentration was within the ideal range of <±15% for all steroids, except for digoxigenin which had a bias of 16.3%. Precision (% RSD) for all CTS was within the ideal range of <15%, except for cinobufotalin, MBG, and TBG at 1.5 ng/mL with 15.5%, 15.5%, and 20%, respectively **(Table 2)**. For beta-blocker bisoprolol precision at 0.3 ng/mL was 15.5%. These were all within our acceptance criteria.

Intra- and inter-assay precision, calculated as a percentage relative standard deviation, was assessed for both serum and plasma. Most data were within desirable limits for a quantitative assay <±15%. Serum intra-assay variability for the 11 CTS averaged 8.0% (range 4.6-15.6%). Bufalin had a variability of 15.6% and beta-blocker bisoprolol 23.6%. Inter-assay variability for CTS averaged 8.4% (range 6.2 – 14.4%) with bisoprolol variation of 20.5%. Plasma intra-assay variability averaged 9.2%, (range 5.9- 17.3%), with only bufalin showing a variability greater than the acceptable range of <15% at 17.3%. Bisoprolol was also above the desirable range at 24.5%. Inter-assay variability in plasma averaged 10.5%, (range 6.3-18.4%), with DHO and ouabagenin over 15% at 16.0% and 18.4%, respectively. Bisoprolol was greater than the acceptable range at 22.2% **(Table 3)**. Bufalin intra-assay variation in serum and inter-assay variation of DHO and ouabagenin in plasma demonstrated reproducible data with variation outside the ideal range but within acceptable criteria. This should be noted when quantifying these steroids in biological samples. The beta-blocker bisoprolol showed the greatest variation, with precision and matrix effects in serum and plasma recording variation above the acceptable limits, therefore this method is only semi-quantitative for bisoprolol.

**Table 3:** Intra and inter-assay precision, matrix effects, recovery and carry-over in serum and plasma for 11 CTS and bisoprolol. Intra and Inter-assay precision, matrix effects and recovery in serum and plasma and percentage carry over at 25 and 250 ng/mL. 500 µl of pooled serum and plasma samples were extracted 10 times on day 1 to assess intra-assay precision and an additional 10 times on another day to assess inter-assay precision. Serum and plasma matrix effects and recovery were calculated following extractions from six samples. Carryover experiments were conducted in triplicate at 25 and 250 ng/mL.

| Steroid | Assay Precision (%CV) |  |  |  | Matrix Effects and Recovery (%) |  |  |  | Carry- over (%) |  |
| --- | --- | --- | --- | --- | --- | --- | --- | --- | --- | --- |
|  | Serum |  | Plasma |  | Serum |  | Plasma |  |  |  |
|  | Intra | Inter | Intra | Inter | Matrix Effects | Recovery | Matrix Effects | Recovery | 25 ng/ml | 250 ng/ml |
| Ouabagenin | 9.9 | 10.4 | 10.7 | 18.4 | 7.6 | 40.0 | -5.1 | 27.4 | 0.001 | 0.001 |
| Dihydroouabain | 6.4 | 7.1 | 7.2 | 16.0 | -2.5 | 70.1 | -14.9 | 68.0 | 0.004 | 0.001 |
| Ouabain | 10.2 | 8.6 | 6.6 | 7.5 | -0.4 | 65.7 | -5.0 | 65.7 | 0.001 | 0.001 |
| Digoxigenin | 4.6 | 7.8 | 6.7 | 8.3 | -14.6 | 76.0 | -16.0 | 83.9 | 0.000 | 0.004 |
| *Bisoprolol | 23.6 | 20.5 | 24.5 | 22.2 | 6.7 | 98.6 | 39.1 | 60.5 | 0.071 | 0.063 |
| Marinobufagenin | 7.7 | 7.1 | 8.5 | 7.0 | -5.2 | 77.4 | 0.7 | 73.0 | 0.027 | 0.020 |
| Cinobufotalin | 6.4 | 6.2 | 6.4 | 8.6 | -5.1 | 83.4 | -5.4 | 81.3 | 0.028 | 0.019 |
| Digitoxigenin | 4.9 | 7.1 | 9.9 | 10.8 | -4.9 | 63.9 | -0.7 | 68.2 | 0.010 | 0.027 |
| Telocinobufagin | 10.8 | 9.2 | 11.8 | 10.5 | -8.5 | 65.6 | -27.6 | 76.4 | 0.000 | 0.005 |
| Digoxin | 7.0 | 7.7 | 9.7 | 8.5 | 2.6 | 78.2 | 0.4 | 79.9 | 0.012 | 0.009 |
| Bufalin | 15.6 | 14.4 | 17.3 | 13.7 | -22.9 | 45.6 | -7.1 | 52.6 | 0.053 | 0.021 |
| Cinobufagin | 4.7 | 7.8 | 5.9 | 6.3 | -11.4 | 71.5 | -1.5 | 77.0 | 0.094 | 0.040 |

\* Not a cardiotonic steroid, included in assay as often prescribed for cardiac related conditions such as AF

#### 3.2.3 Matrix Effects and Recovery

Recovery was optimised for all analytes, which was highly challenging due to the variability in polarity of the CTS. In serum, average recovery was 67%, (range 40.0-83.4%), and in plasma 68.4%, (range 27.1 to 83.9%). Ouabagenin recovery was the lowest, with recovery from serum and plasma being 40.0 and 27.4%, respectively. Bufalin recovery was low in both serum and plasma, 45.6% and 52.6%, respectively **(Table 3)**. Although precision data were within acceptable ranges, caution should be used when quantifying ouabagenin and bufalin.

Matrix effects in serum were in the desirable range (<±15%), except for bufalin (-22.9%). Matrix effects in plasma were generally higher with 9/11 CTS reaching the desirable levels of <±15%. Digoxigenin was acceptable at -16.0%, however, TBG was outside our acceptable criteria at -27.6%. Therefore, quantification of CTS in serum may be preferable to plasma due to the lower matrix effects observed **(Table 3)**.

#### 3.2.4 Carryover

Carryover was assessed at two concentrations, the ULOQ, (25 ng/mL), and ten times this concentration (250 ng/mL). Carryover ranged from 0.0001% to 0.094%, for both concentrations, all within our acceptability criteria of <2% **(Table 3)**.

#### 3.2.5 Bisoprolol cross-reactivity

Samples were spiked with bisoprolol to replicate patients who receive therapy with β-blockers and digoxin. The retention time for bisoprolol was 5.8 minutes. Due to its unique retention time and mass transitions, it does not cross-react with any of the CTS. If a method is required for investigating bisoprolol independently of CTS our method would be a promising starting point. We were able to quantify this analyte between 0.025 and 25 ng/mL, with minimal matrix effects in serum, and reproducible recoveries of 98.6% (serum) and 68.2% (plasma) **(Table 3)**. However, additional work would be required to reduce matrix effects and improve intra and inter-assay precision, as these were outside the acceptable range in both serum and plasma.

#### 3.2.6 Stability

Stability was assessed at three concentrations (0.3, 1.5, and 15 ng/ml). Different aliquots were extracted eight times over a three-month period. Accuracy expressed as percentage bias from theoretical concentrations was determined in all experiments. Average bias was 6.3% (-17.5 to 15%), 14.3% (-15.2 to -23.1%), and 0.6% (-4.2 to 11.1%) at low, medium, and high concentrations, respectively. Precision measured as percentage variance over all eight extractions was an average (range) of 12.5% (7.3 to 17.7%), 10.6% (5.1 to 18.1%), and 8.8% (3.7 to 17.2%) at low, medium, and high concentrations, respectively. These data demonstrates that all the CTS were stable under the storage conditions used, without significant impact on quantitation.

### 4.0 Discussion

Here we describe a liquid chromatography tandem-mass spectrometry method for quantification of the cardenolides ouabain, dihydroouabain (DHO), ouabagenin, digoxigenin, and digoxin, and bufadienolides bufalin, cinobufotalin, cinobufagin, marinobufagenin (MBG) and telocinobufagin (TBG). To our knowledge, no reported assay measures more than three CTS in a single method or includes CTS from both the cardenolide and bufadienolide classes. Our method enables the study of the interaction between these ten CTS and digoxin and their effects on heart rate and cardiac function.

Multiple studies have detected CTS in serum or plasma. In general, endogenous CTS were measured using ELISA, though, several mass spectrometry-based methods were employed to measure ouabain [25], digoxin [25], MBG [29], TBG, [27], and bufalin [30]. Our method was developed to quantify multiple CTS in serum and plasma patient samples in the future. Serum and plasma were used in this study to determine and compare matrix effects and extraction reproducibility of CTS spiked biological samples.

Validation of the method when applied to serum or plasma was completed to describe the robustness of the method, including the quantification range, reproducibility, and reliability of the data. Although structurally similar, CTS demonstrate vast differences in ionisation efficiency and polarity development of a single extraction and analytical method challenging. Therefore, a Pareto-optimal solution was developed. A Pareto-optimal method demonstrates acceptable recovery, reproducibility, accuracy, and precision for all analytes, but is not optimised for a single analyte. Producing a useful, research tool for investigation into the function and metabolism of CTS.

Acceptable validation criteria in serum were reached for DHO, ouabain, digoxigenin, digoxin, cinobufotalin, cinobufagin, MBG, TBG and digitoxigenin. For plasma, acceptable validation criteria were met for DHO, ouabain, digoxigenin, digoxin, cinobufotalin, cinobufagin, MBG, and digitoxigenin. Analytes that did not meet the acceptable quantification criteria were ouabagenin and bufalin in serum and plasma, and TBG in plasma. Measurement of ouabagenin by this method was semi-quantitative as recovery was below acceptable limits. Ouabagenin and bufalin quantitation could be improved using a more suitable internal standard to account for the poor recovery, or via further development of the extraction procedure. Due to the observed matrix effects, bufalin and TBG in plasma was qualitative. Factors that affect the ability of these compounds to reach the acceptability criteria are poor recovery or matrix effects causing variance in the data. Further optimisation of the extraction method, the inclusion of suitable internal standards, and optimisation of the chromatography methods could improve recovery and reduce matrix effects. However, this should be completed with caution, as improvement in the recovery/matrix effects of a single analyte is often detrimental to the recovery or cross-reactivity of another.

In this method, we extracted CTS and bisoprolol from 500 μL of serum/plasma with the final reconstitution volume of 200 μL. At this ratio, we were able to quantify added CTS, while keeping the matrix effects acceptably low. We could theoretically improve sensitivity, by injecting a more concentrated sample onto the LC-MS/MS. This could be achieved in three ways 1. increasing the volume of sample, although this is limited by the bed volume of the SPE cartridges, or 2. reconstituting our samples in a lower volume, or 3. injecting more sample onto the LC-MS/MS. However, all the improvements in absolute signal (peak area) would be counterbalanced by increases in matrix effects, possibly resulting in a higher LLOQ. Our method presents a solution that considers all these factors.

The calibration ranges we employed here were based on literature searches quantifying CTS either by ELISA or mass spectrometry. There are a variety of sample volumes, extraction methods, and analysis procedures that make comparison of our LLOQ to other methods difficult. For example, Komiyama and colleagues identified ouabain-like factor and digitalis-like factor from 80 mL of plasma [25]. This large sample volume makes their method not viable for routine analysis, with the possibility of high matrix effects. Kirby et al [45] developed an LC-MS/MS method for digoxin quantification extracting from 1 mL of plasma and quoted a LLOQ of 0.05 ng/mL, this is double our quoted LLOQ of 0.025 ng/mL from 500 μL of sample. Additionally, their method did not quantify based on a Li-adduct, possibly explaining the difference in the LLOQ. Komiyama and colleagues [27] detected TBG and MBG (again without the use of Li-adducts) in 1 mL of plasma using high-resolution mass spectrometry quoting a LOQ of 1 ng/mL for both compared to 0.05 and 0.025 ng/mL described here. The most similar method in the literature is from Baecher [31] and colleagues, who formed Li-adducts and conducted a similar extraction methodology to us. They only investigated ouabain, allowing them to achieve optimal sensitivity of this single CTS and a LLOQ of 1.7 pmol/L, which is more sensitive than we were able to achieve in our profiling method. Interestingly, they did not detect any endogenous CTS by their method and concluded that immunoassays do not detect ouabain, but due to cross-reactivity there was a false positive.

Plasma demonstrated greater variability in inter and intra-assay precision than serum. Matrix effects varied between plasma and serum with larger differences observed with bisoprolol and TBG. Digoxigenin and digitoxigenin had intermittent matrix effects in the quantifier but not the qualifier ions (see **Supplementary Figures 2D, 3H, and 4H**). In the serum (Sigma Aldrich) and plasma (six healthy volunteers) investigated here, there were no detectable endogenous CTS using our method.

Serum and plasma have many similarities, both are complex matrices derived from blood, which is a mixture composed of water, albumin, globulins, hormones, enzymes, amino acids, and more. In addition, there can be significant biological variation between patients, with gender, age, time of day, diet, hydration, exercise, stress, and disease status all altering the composition of blood, leading to different matrix effects from person to person. Plasma is inherently more complex than serum with more chemicals present, and this is likely why we observed higher matrix effects in plasma than serum. Plasma collection tubes also contain an anticoagulant such as EDTA which prevents clotting and could lead to additional matrix effects. Other considerations when exploring matrix effects include tube type e.g., clotting factor used, plastics, and additives as well as matrix contaminants being introduced during the extraction and analytical methods themselves. These factors may impact analyte recovery as well as contribute to matrix effects if not removed. In research, it is best practice to use the same tube type and storage conditions to minimise variation. The optimal collection tube type to be used with our method is yet to be explored.

### 4.1 Limitations and Improvements

Where recovery is below 60%, (two CTS in serum and plasma), additional sample preparation steps may improve this. Alternatively, concentrating the sample post-extraction can improve mass spectrometry sensitivity. This should be attempted with caution as concentrating samples can increase matrix effects.

Matching stable isotope-labelled internal standards are considered a prerequisite to appropriately control for matrix effects and extraction losses and to allow for accurate and precise quantification [46,47]. A limitation of our study is stable isotope-labelled internal standards for all CTS were not available, and a surrogate had to be used. The inclusion of isotopically labelled internal standards identical to the analyte of interest should improve the accuracy and precision data as well as compensate for lower recovery. However, our validation data demonstrated that the use of the surrogate internal standards were adequate as acceptable validation criteria were met. This reduces the cost of the assay as the customised synthesis of deuterium- or ^13^C-labelled internal standards are expensive.

We investigated 11 CTS in this method, however, there could be additional cross-reactivity from other CTS and/or other steroidal analytes in this method. We minimised the chances of this through use of lithium adducts, optimisation of solid phase extraction, CORTECS column chemistry, and a novel LC gradient.

In these experiments, we used PBS 0.1%BSA as a surrogate matrix. This matrix ratio has been used widely for steroid quantification in biological samples [42, 48–51]. Liu and colleagues investigated different surrogate matrixes including PBS, twice charcoal-stripped serum, NaCl, and water to investigate the quantification of five steroid hormones. They demonstrated no significant differences in their data depending on the matrix used [52]. CTS- stripped serum or plasma may be a better representation of the matrix, however, to our knowledge there is no certified CTS-stripped serum available. Additionally, charcoal stripping of serum removes many additional analytes producing a variable surrogate matrix. An additional advantage of using a PBS based surrogate matrix is no batch-to-batch variation. Therefore PBS 0.1%BSA is a suitable surrogate matrix for the preparation of calibration series and QC samples to quantify CTS.

Finally, we were only able to determine matrix effects at one spiked concentration due to limited sample availability. As there was no detectable CTS in our healthy serum or plasma samples, we were unable to discern matrix effects at endogenous concentrations.

### 5.0 Conclusion

UHPLC-MS/MS with post-column infusion of lithium chloride enables profiling of 11 cardiotonic steroids. This method was analytically assessed for application to human serum and plasma, with acceptable validation criteria being reached for most parameters. This method can be applied to biological samples to aid studies where endogenous CTS measurements are required and to investigate the role of CTS in disease. The method can also be used to aid studies examining stratified therapeutic approaches in patients with heart failure, atrial fibrillation, hypertension, and renal disease.

## Acknowledgments

The authors thank all volunteers who donated blood samples and Prof M Viant and Dr J Bradbury, both University of Birmingham, UK, for the use of MUSCLE software.

## Abbreviations

AF: Atrial fibrillation
BSA: Bovine serum albumin
CTS: Cardiotonic steroids
DHO: Dihydroouabain
ELISA: Enzyme-linked immunoassay
LC-MS/MS: Liquid chromatography tandem-mass spectrometry
LLOQ: Lower limit of quantification
MBG: Marinobufagenin
MUSCLE: Multi-objective Unbiased optimisation of Spectrometry via closed Loop Experimentation
QC: Quality control
RSD: Relative standard deviation
SPE: Solid phase extraction
TBG: Telocinobufagin
UHPLC: Ultra-high performance liquid chromatography
ULOQ: Upper limit of quantification

