## Supplemental files for "A cardiotonic steroid multiplex method using ultra-high-performance liquid chromatography-tandem mass spectrometry"

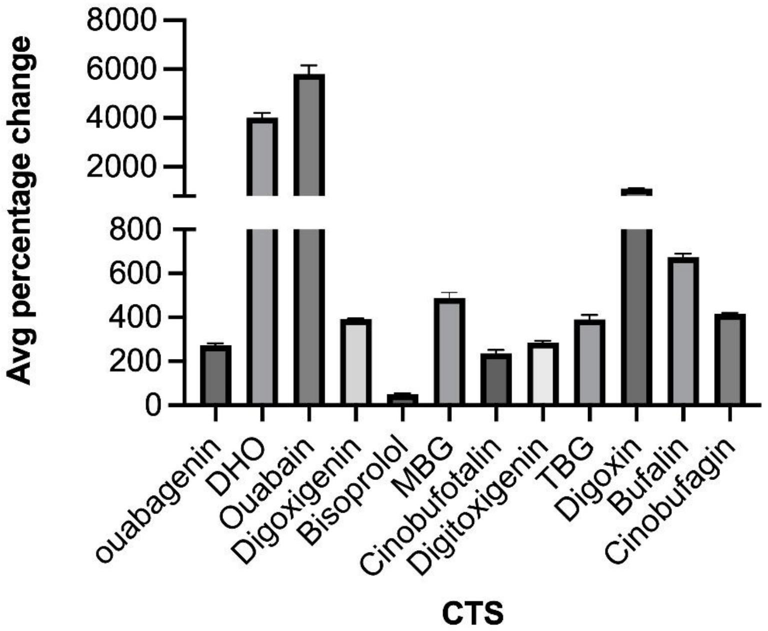


**Supplementary Figure 1.** Percentage increase in absolute peak area comparing the protonated molecular ion to the lithium adduct. Data is based on three injections of the same CTS mix (50ng/mL) without and three injections with LiCl post-column infusion. Bisoprolol does not form a lithium adduct.

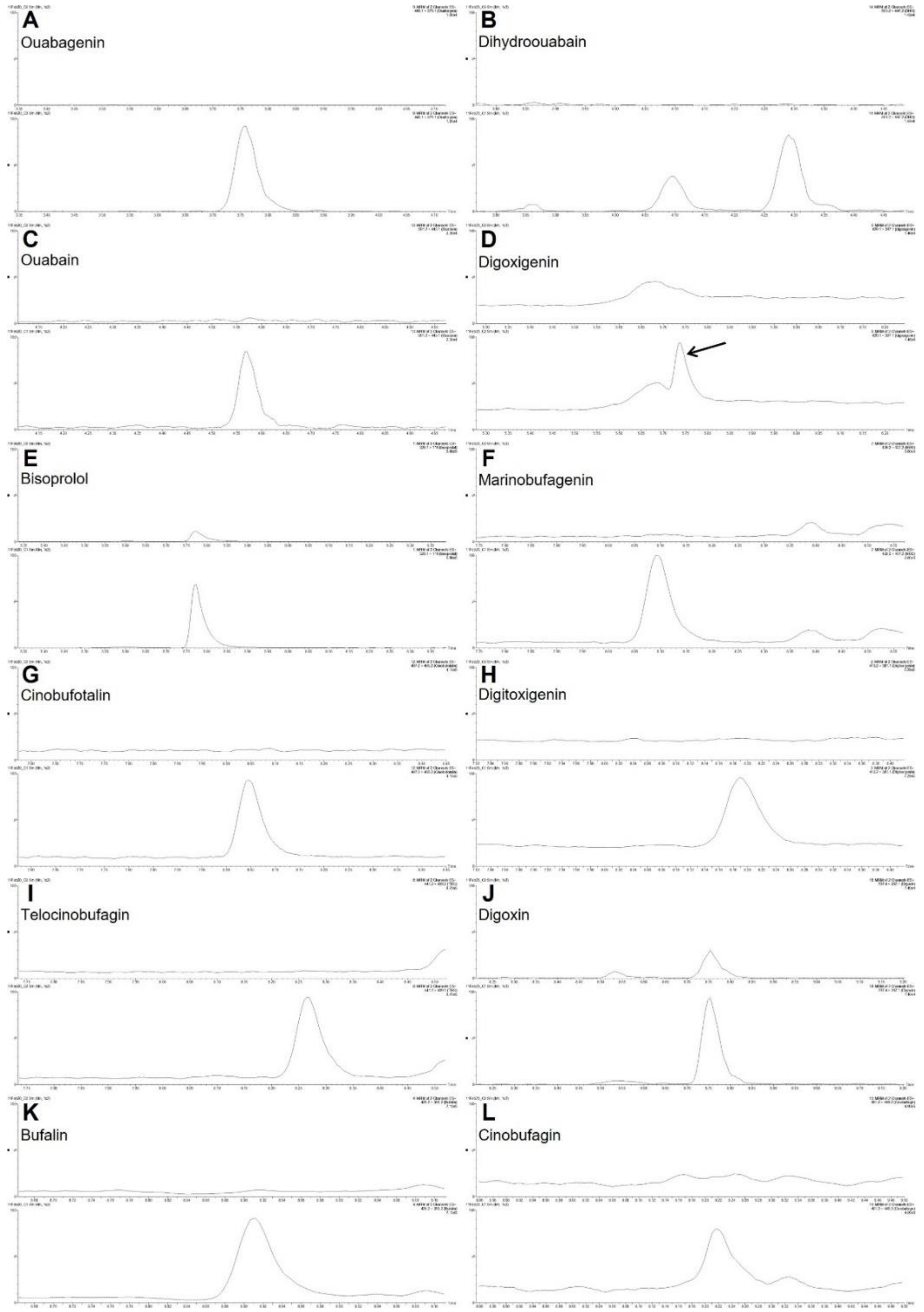


**Supplementary Figure 2.** Chromatograms for the quantifier mass transitions (MRMs) for each cardiotonic steroid and bisoprolol at the calculated lower limit of quantification. The upper chromatogram for each analyte is 0 ng/ml (calibration blank with added internal standards) and the lower chromatogram is the peak at the lower limit of quantification for that analyte. The peak intensity is relative, and they have been smoothed by MassLynx software 1 x 2. (A) Ouabagenin and 0.1 ng/ml, (B) DHO and 0.025 ng/ml, note DHO standard forms two peaks and only the second peak was used (C) Ouabain and 0.025 ng/ml, (D) Digoxigenin and 0.05 ng/ml, the arrow points to the correct peak (E) Bisoprolol and 0.025 ng/ml, (F) MBG and 0.025 ng/ml, (G) Cinobufotalin and 0.025 ng/ml, (H) Digitoxigenin and 0.025 ng/ml, (I) TBG and 0.05 ng/ml, (J) Digoxin and 0.025 ng/ml, (K) Bufalin and 0.025 ng/ml, and (L) Cinobufagin and 0.025 ng/ml.

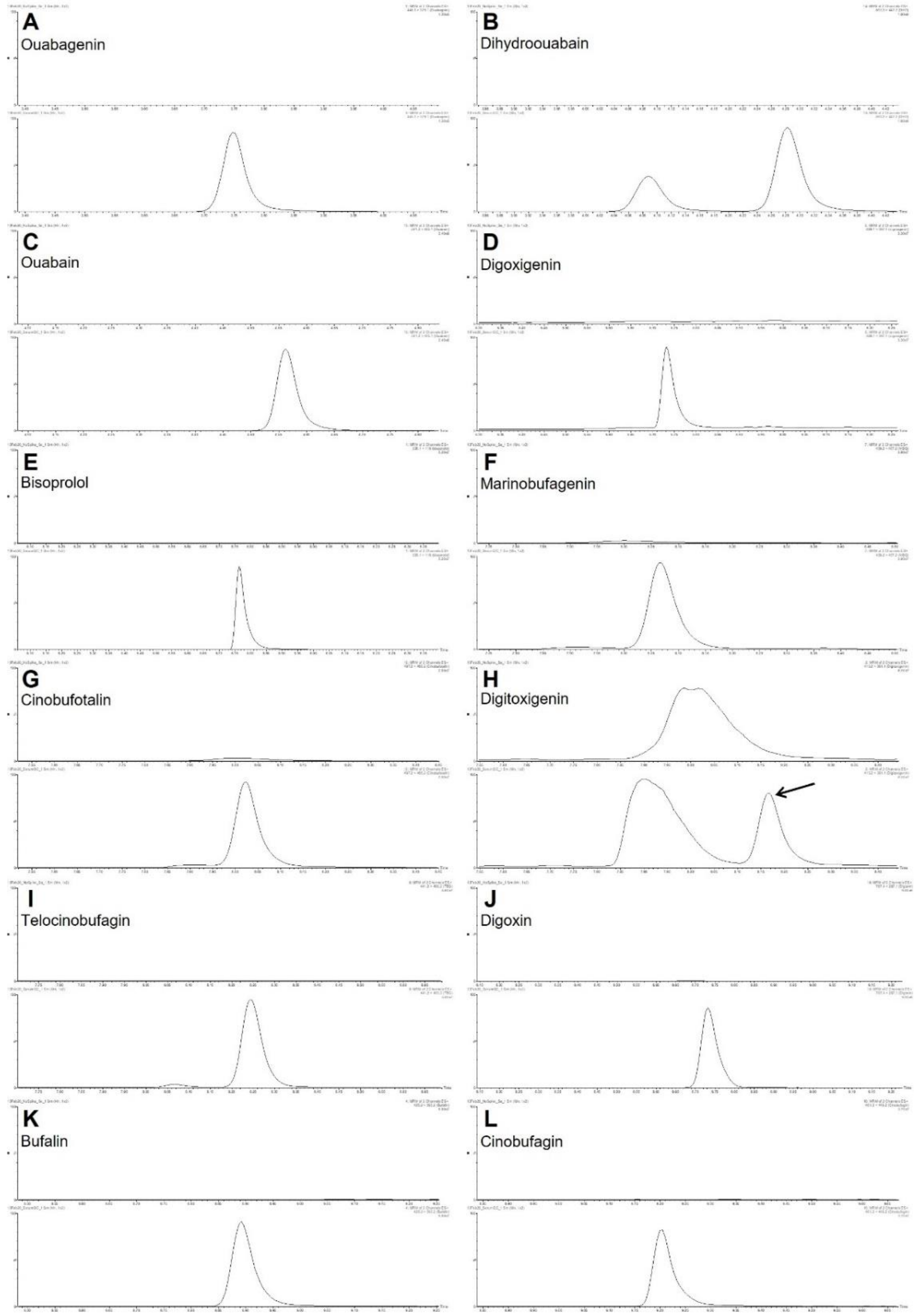


**Supplementary Figure 3.** Chromatograms for the quantifier mass transitions for each cardiotonic steroid and bisoprolol in pooled serum. The upper chromatogram is an example of pooled serum (500 µl extracted) with only internal standards added and the lower chromatogram is with the pre-extraction addition of a 2 ng/ml mix of all CTS and bisoprolol. The peak intensity is relative, and they have been smoothed by MassLynx software 1 x 2. (A) Ouabagenin (B) DHO (two peaks with only the second peak used in analysis), (C) Ouabain, (D) Digoxigenin, (E) Bisoprolol, (F) MBG, (G) Cinobufotalin, (H) Digitoxigenin, with an arrow pointing to correct peak (I) TBG, (J) Digoxin, (K) Bufalin, and (L) Cinobufagin.

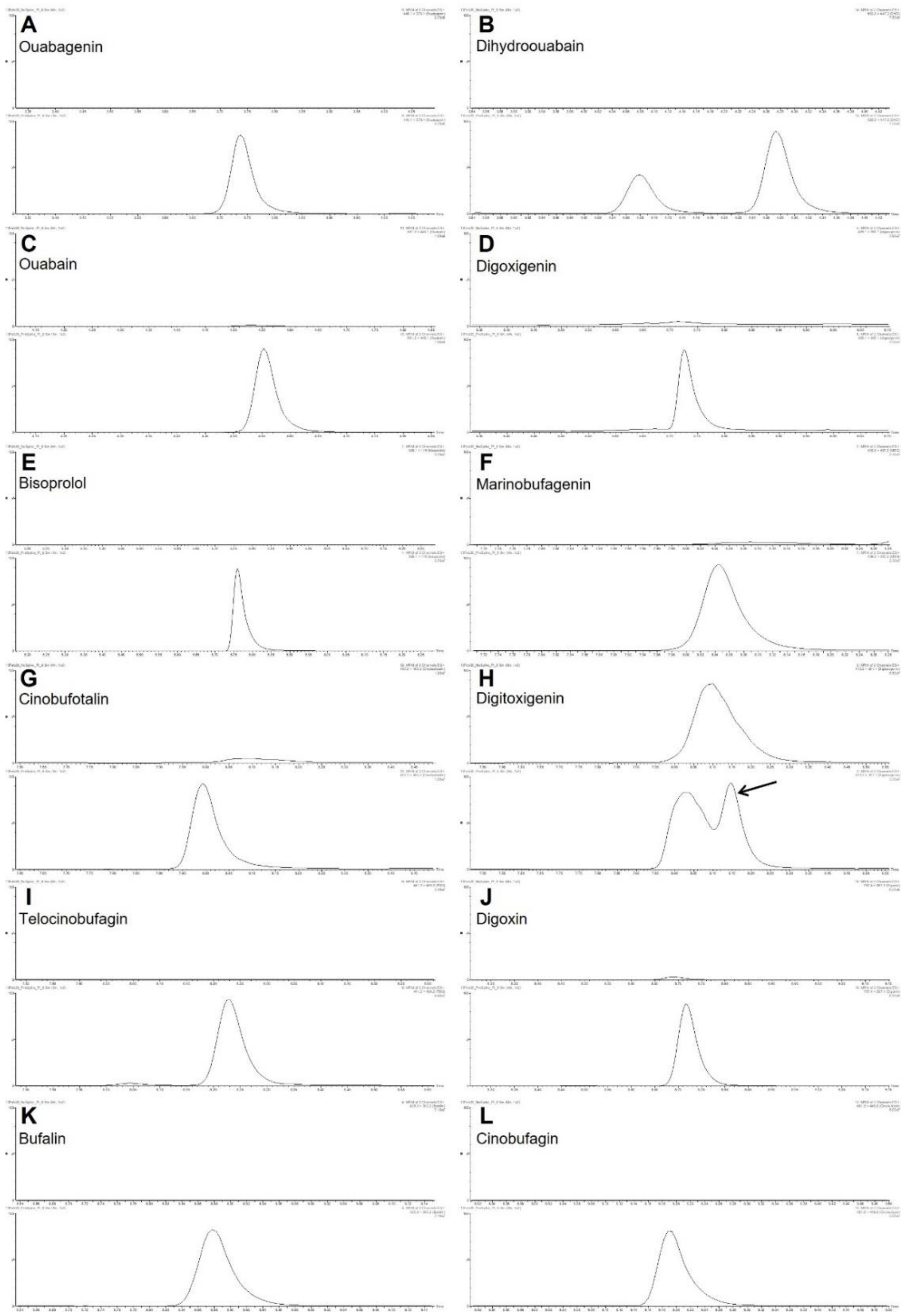


**Supplementary Figure 4.** Chromatograms for the quantifier mass transitions for each cardiotonic steroid and bisoprolol in a plasma sample. The upper chromatogram is an example of plasma (500 µl extracted) with only internal standards added and the lower chromatogram is with the pre-extraction addition of a 2 ng/ml mix of all CTS and bisoprolol. The peak intensity is relative, and they have been smoothed by MassLynx software 1 x 2. (A) Ouabagenin (B) DHO (two peaks with only the second peak used in analysis), (C) Ouabain, (D) Digoxigenin, (E) Bisoprolol, (F) MBG, (G) Cinobufotalin, (H) Digitoxigenin, with an arrow pointing to correct peak (I) TBG, (J) Digoxin, (K) Bufalin, and (L) Cinobufagin.

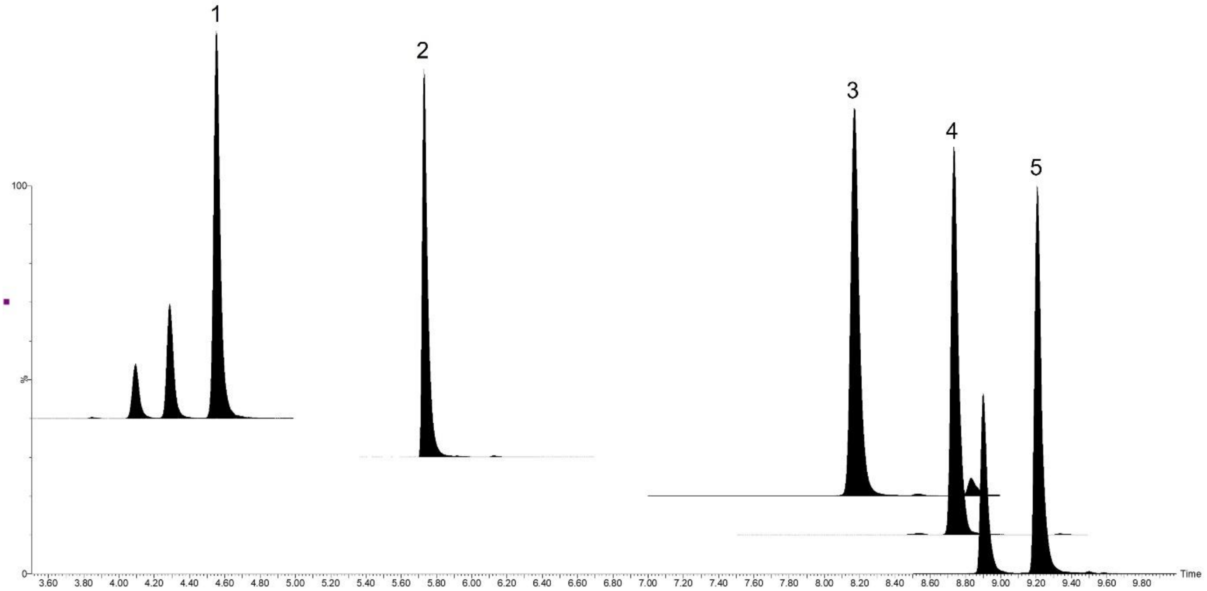


**Supplementary Figure 5.** Chromatograms for the quantifier mass transitions for the cardiotonic steroid deuterated internal standards. Peaks are (1) ouabain-d3, (2) digoxigenin-d3, (3) digitoxigenin-d3, (4) digoxin-d3, and (5) cinobufagine-d3. The peak intensity is normalised to the largest peak for each internal standard and they have been smoothed by MassLynx software 1 x 2.
